# A Minimally Destructive Protocol for DNA Extraction from Ancient Teeth

**DOI:** 10.1101/2020.08.19.256412

**Authors:** Éadaoin Harney, Olivia Cheronet, Daniel M. Fernandes, Kendra Sirak, Matthew Mah, Rebecca Bernardos, Nicole Adamski, Nasreen Broomandkhoshbacht, Kimberly Callan, Ann Marie Lawson, Jonas Oppenheimer, Kristin Stewardson, Fatma Zalzala, Alexandra Anders, Francesca Candilio, Mihai Constantinescu, Alfredo Coppa, Ion Ciobanu, János Dani, Zsolt Gallina, Francesco Genchi, Emese Gyöngyvér Nagy, Tamás Hajdu, Magdolna Hellebrandt, Antónia Horváth, Ágnes Király, Krisztián Kiss, Barbara Kolozsi, Péter Kovács, Kitti Köhler, Michaela Lucci, Ildikó Pap, Sergiu Popovici, Pál Raczky, Angela Simalcsik, Tamás Szeniczey, Sergey Vasilyev, Cristian Virag, Nadin Rohland, David Reich, Ron Pinhasi

**Author notes:** These authors contributed equally (Éadaoin Harney, Olivia Cheronet). Correspondence to Ron Pinhasi (Tel). Correspondence may also be addressed to Éadaoin Harney or Olivia Cheronet.

## Abstract

Ancient DNA sampling methods—although optimized for efficient DNA extraction—are destructive, relying on drilling or cutting and powdering (parts of) bones and teeth. As the field of ancient DNA has grown, so have concerns about the impact of destructive sampling of the skeletal remains from which ancient DNA is obtained. Due to a particularly high concentration of endogenous DNA, the cementum of tooth roots is often targeted for ancient DNA sampling, but standard destructive sampling methods often result in the loss of at least one entire root. Here, we present a minimally destructive method for extracting ancient DNA from dental cementum present on the surface of tooth roots. This method does not require destructive drilling or grinding, and, following extraction, the tooth remains safe to handle and suitable for most morphological studies, as well as other biochemical studies, such as radiocarbon dating. We extracted and sequenced ancient DNA from 30 teeth (and 9 corresponding petrous bones) using this minimally destructive extraction method in addition to a typical tooth sampling method. We find that the minimally destructive method can provide ancient DNA that is of comparable quality to extracts produced from teeth that have undergone destructive sampling processes. Further, we find that a rigorous cleaning of the tooth surface combining diluted bleach and UV light irradiation seems sufficient to minimize external contaminants usually removed through the physical removal of a superficial layer when sampling through regular powdering methods.

## INTRODUCTION

Over the past decade, the field of ancient DNA has experienced a rapid increase in the number of ancient genomes published each year (Slatkin and Racimo 2016) as a consequence of advances in ancient DNA sampling (Gamba et al. 2014; Damgaard et al. 2015), extraction (Dabney et al. 2013a; Rohland et al. 2018), and enrichment (Carpenter et al. 2013; Fu et al. 2013) techniques. As our ability to sequence large numbers of ancient individuals has increased, discussions about the destructive nature of ancient DNA sampling—which typically requires drilling or cutting and powdering ancient bones and teeth—have become more prominent (Makarewicz et al. 2017; Prendergast and Sawchuk 2018; Sirak and Sedig 2019). The identification of the osseous inner ear, and specifically the cochlea (located in the petrous portion of the temporal bone), as an optimal source of ancient DNA (Gamba et al. 2014; Pinhasi et al. 2015; Pinhasi et al. 2019) is one of the driving factors in this revolution, making it possible to access ancient DNA from geographic regions with climatic conditions unfavorable to ancient DNA preservation. However, accessing this optimal source of ancient DNA results in the destruction of the inner ear morphology, which is a valuable source of morphological information (de León et al. 2018). While there are protocols that reduce the destructive nature of sampling, by sampling from the ossicles of the inner ear (Sirak et al. 2020) or performing targeted drilling of the cochlea through the cranial base of complete or reconstructed crania (Sirak et al. 2017), some destruction (including that of morphologically-informative inner ear components) is inevitable. As a consequence, this and other less-invasive methods may be considered unsuitable in cases where samples are of particular anthropological value and are subject to stringent restrictions on permissible sampling practices.

Teeth are a valuable alternative to the sampling of the cochlea (Gamba et al. 2014; Damgaard et al. 2015), especially because they are particularly numerous in osteological collections, due to the fact that individuals have many more teeth than petrous bones and to their resistance to taphonomic decomposition. Despite this, little has been published outlining optimal practices for sampling from teeth. Traditionally, the standard practice has been to grind or drill large chunks of the tooth root to a powder (Rohland and Hofreiter 2007), as the crown enamel is largely inorganic and is therefore unlikely to contain a substantial amount of endogenous DNA (Higgins and Austin 2013). In an attempt to minimize potential external contaminants, the surface layer is often removed to access the “untouched” dentine and pulp. However, this practice removes some, if not all, of the thin layer of cementum that coats the inferior portion of dental roots.

The cellular cementum is rich in cementocytes, which are DNA containing cells that remain encased in the mineral structure of the tooth after death (Bosshardt and Selvig 1997). Cementum also shares several histological properties with the cochlear region of the petrous that are thought to contribute to its high level of DNA preservation, including similarities between cementocytes (Zhao et al. 2016) and osteocytes, which are hypothesized to be serve as repositories of ancient DNA in bones (Bell et al. 2008; Pinhasi et al. 2015). Like the cochlea, cementum also does not undergo remodeling (but, unlike the cochlea, it continues to accumulate throughout life) and the haphazard organization of collagen fibers in cementum resembles that of woven bone (Freeman 1994; Grzesik et al. 2000). Assessment of DNA preservation in ancient teeth shows that dental cementum contains a substantially higher proportion of endogenous DNA than dentine from the same tooth (Damgaard et al. 2015). Furthermore, in a direct comparison between cementum and petrous samples, Hansen et al. (2017) find that cementum and petrous yield a comparable amount of endogenous DNA in well-preserved samples, although in poorly-preserved individuals, the petrous yields a higher proportion of endogenous molecules. The only published method for sampling DNA from the cementum recommends a targeted method for extracting DNA from teeth using an “inside-out” approach that involves removing the crown and subsequently using a fine drill to remove as much pulp and dentine as possible from the tooth root to ultimately obtain a “case” of cementum (Damgaard et al. 2015). However, this valuable approach may still not be able to perfectly isolate the extremely thin and brittle layer of cementum, which ranges from 20-50 μm thick at the cementoenamel junction, to 150-200 μm thick at the apex of the root (Freeman 1994).

Here, we present an alternative, minimally destructive protocol for sampling ancient DNA from tooth cementum that does not require drilling or cutting, thereby maintaining the morphological integrity of the tooth. The technique isolates ancient DNA from the cementum of tooth roots by directly exposing the outermost layer of a portion of the tooth root to a lysis buffer for a short incubation period, following a non-destructive decontamination procedure. Similar less destructive methods have been reported in previous PCR-based mitochondrial ancient DNA studies (Rohland et al. 2004; Bolnick et al. 2012) and in forensic contexts (Correa et al. 2019). However, the ancient DNA obtained using these strategies was typically less well preserved and of a lesser quantity than DNA obtained using more destructive methods. Additionally, in some cases (Rohland et al. 2004), the hazardous chemicals used during sampling may have compromised safe handling and future chemical analyses of the remains. In this study, we conduct a systematic evaluation of the application of a minimally destructive sampling technique in a next generation sequencing context. This protocol is further optimized by enabling targeted sampling from the very thin dental cementum layer, which increases the quality of ancient DNA sampled from the tooth while giving researchers the ability to fully preserve the dental crowns and all but the fine external detail of the roots. After sampling, teeth can be safely handled and remain suitable for subsequent morphological and biomolecular analyses, such as radiocarbon dating (Korlević et al. 2018).

## RESULTS

We selected thirty ancient individuals (Table 1; Supplementary Table 1) for a comparative analysis of the quality of ancient DNA—as measured through metrics such as the proportion of endogenous molecules of shotgun data, sample complexity and contamination rate—that could be obtained from an individual using this minimally destructive extraction method versus standard sampling procedures that rely on cutting and powdering tooth samples. From each individual we sampled a single multi-rooted tooth, from which the roots were removed via cutting (note that the tooth roots were cut in order to make it possible to process the samples using several independent methods, but cutting is not required by the minimally destructive sampling protocol) and were each randomly assigned to undergo one of the following extraction treatments. We extracted ancient DNA from a tooth root that was processed using the minimally destructive extraction protocol described in this paper (Method “MDE”; for “Minimally Destructive Extraction”) and a second whole tooth root of the same tooth, that was completely powdered via milling (Method “WTR”; for “Whole Tooth Root”). We also generated extracts from powder produced from petrous bones for 10 of the same individuals using the method described by Pinhasi et al. (2019) (Method “P”; for “Petrous”). In one case (individual 3), we discovered through subsequent bioinformatic analyses that the petrous bone and tooth sampled did not originate from the same individual, and we therefore exclude the petrous bone results from further analyses. DNA preservation in two individuals (5 and 6) was uniformly poor, with no more than 10,000 sequences aligning to the 1.24 million sites captured through targeted enrichment (out of ~5 million unique reads sequenced) from any of the libraries generated. Furthermore, all of these double-stranded libraries exhibited C-to-T damage rates at the terminal ends of molecules of less than 3%—the recommended minimum threshold for assessing ancient DNA authenticity in partially UDG treated libraries (Rohland et al. 2015). These samples are considered to have ‘failed’ screening for authentic ancient DNA and are not included in the statistical analyses. Additionally, individual 22 yielded relatively poor results for both treatments. Only 533 reads (out of ~4 million unique reads sequenced) aligned to the 1.24 million sites targeted in the nuclear genome for the MDE treatment, making it impossible to calculate several of the reported metrics. While we did obtain enough reads (23,239 reads out of ~18 million unique reads sequenced) for some analyses to produce results for the tooth root that underwent Method WTR, the relatively low rate of mitochondrial match to the consensus (0.860) suggests that this sample is likely contaminated. Based on these results, we also chose to exclude individual 22 from statistical analyses. However, we note that there are no significant changes to the reported statistics when the excluded individuals are included in calculations for which metrics from both treatments are available (Supplementary Table 2). For all statistical calculations, we included data from all other samples, which were processed as either double-stranded (samples 1-10) or single-stranded (samples 11-30) libraries. Results where each of these methods were analyzed separately are reported in Supplementary Table 2.

**Table 1.**
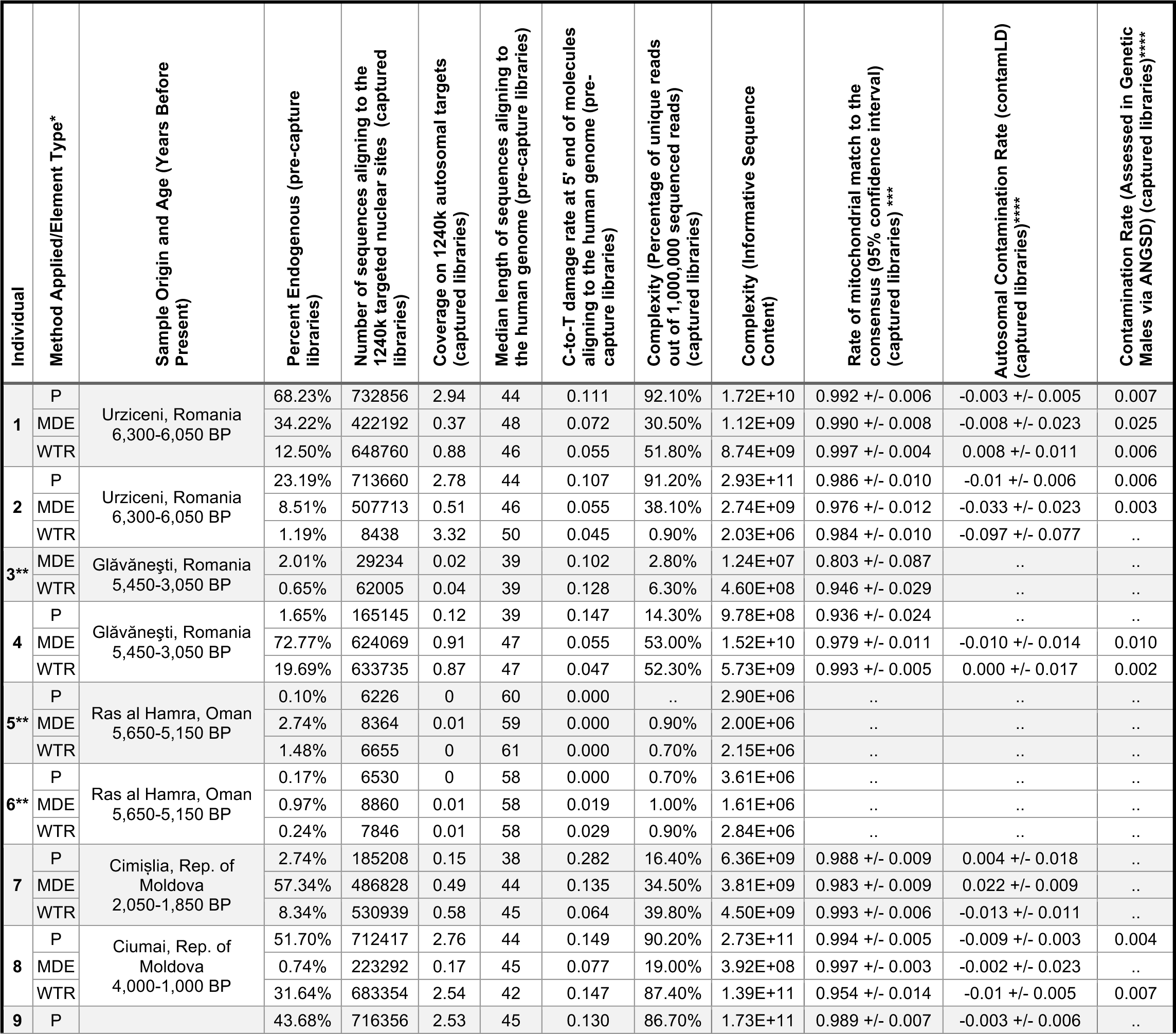
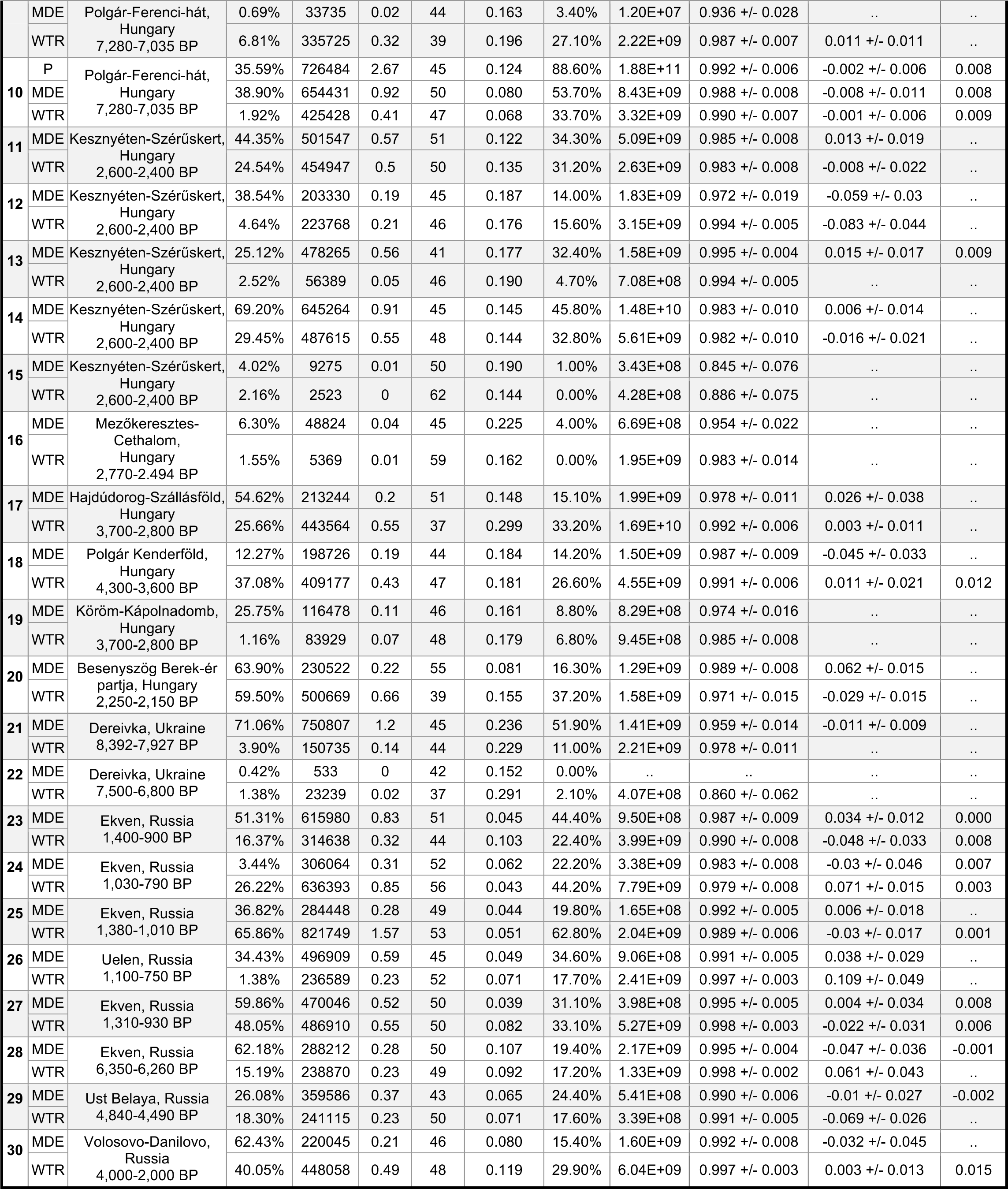
Sample Information. All estimates are made based on data produced from libraries that underwent the 1240k capture unless otherwise specified. *Sampling/Extraction Methods: P-Powdered Petrous Bone (Pinhasi et al. 2019), standard extraction (Dabney et al. 2013a); MDE-Tooth Root processed via Minimally Destructive Extraction; WTR-Whole Tooth Root, powdered with standard extraction (Dabney et al. 2013a). Extracts for individuals 1-10 were processed entirely manually and underwent partial UDG treatment followed by double stranded library preparation, while extracts for individuals 11-30 were processed robotically following incubation in extraction buffer (Rohland et al 2019, buffer D) and processed using USER treatment followed by single stranded library preparation. ** Note that sample 3P was excluded from comparisons as it was determined bioinformatically that the petrous bone and tooth sampled did not originate from the same individual. Note also that the DNA preservation in samples 5 and 6 was too poor for further analysis. *** Contamination estimates are not reported for samples which did not produce sufficient quality data to generate a contamination estimate based on either mitochondrial, autosomal or X-chromosome data. For X-chromosome based contamination estimates, ANGSD can only estimate contamination rates for individuals determined to be genetically male. Individuals who are female or for whom sex cannot be determined (sex ND) are noted.

### Physical Impact of Minimally Destructive Extraction Protocol

We photographed each tooth root processed using the minimally destructive extraction protocol immediately prior to extraction and 24 hours after extraction to allow for the complete drying of the roots (Figure 1; Supplementary Figure 1). A slight degradation of the outer tooth root surface is visible for many of the samples, as the portion of the tooth root exposed to extraction buffer shows a visible change in color and/or diameter relative to the unexposed portion. In the case of two of the most poorly preserved samples (individuals 5 and 6), the tooth roots—one of which broke in two when cut from the tooth crown—crumbled during removal of the parafilm that covered the tops of the roots after the incubation in extraction buffer. These results suggest that users should exercise caution when applying this method to very friable teeth that are already susceptible to crumbling or being crushed.

**Figure 1.**
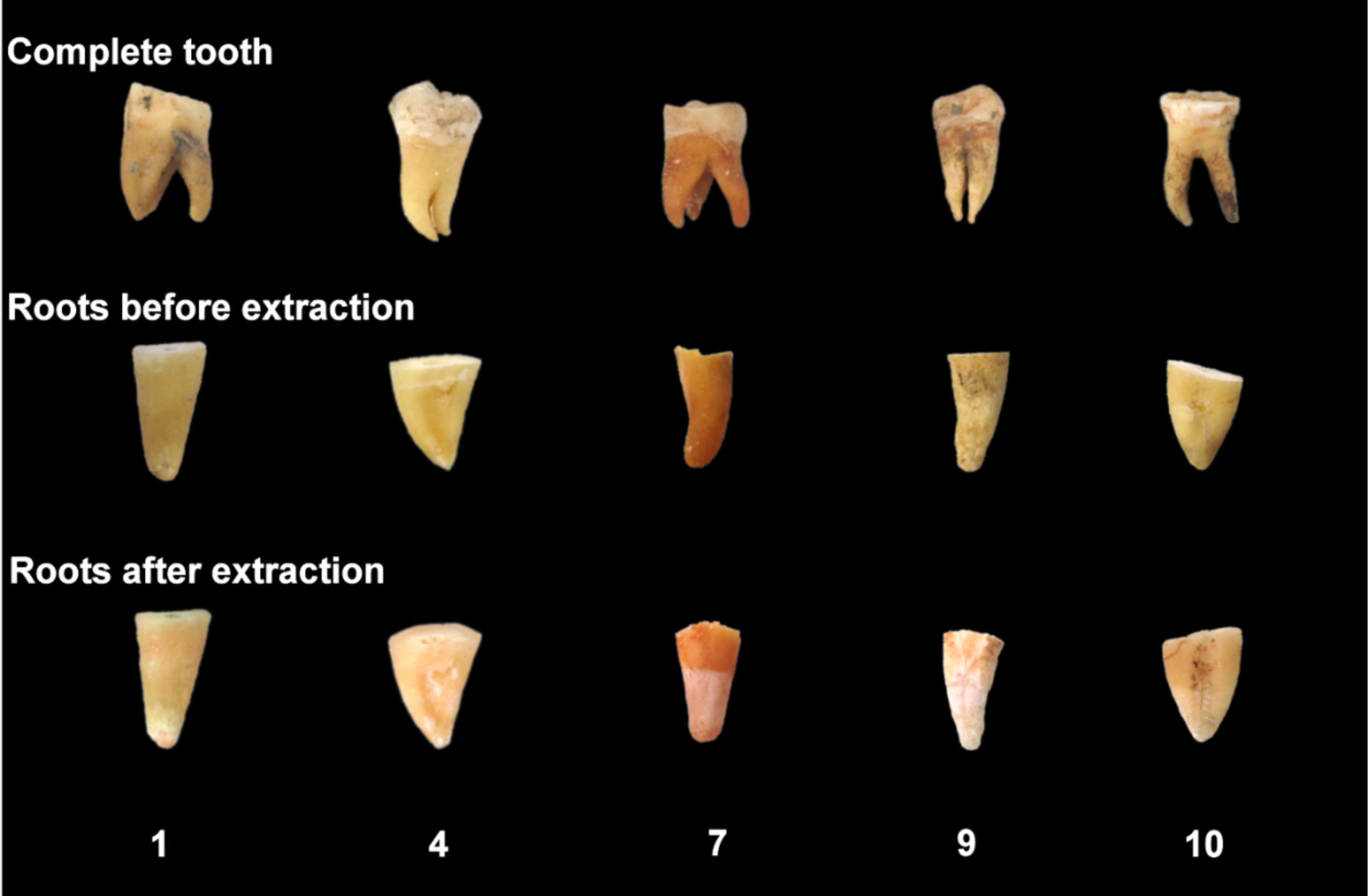
Tooth roots before and after minimally destructive extraction. The complete tooth is shown prior to processing (top). Tooth roots are shown immediately prior to extraction (middle) and 24 hours after extraction (bottom). See Supplementary Figure 1 for before and after images of all sampled teeth.

### Comparison of Minimally Destructive Extraction Protocol versus Powder-Based Extraction Protocols

Following bioinformatic processing, we generated summary statistics for each extract, including metrics of sample complexity and contamination rates (Table 1, Supplementary Table 1). In the following section, for each individual we compare the quality of ancient DNA yielded by the minimally destructive extraction method (Method MDE) to that produced by the destructive, traditional sampling methods (Methods WTR and P), using a Wilcoxon signed-rank test. The null hypothesis is that the difference between pairs of data generated using Method MDE and Method WTR or P follows a symmetric distribution around zero. The alternative hypothesis is that the difference between the paired data does not follow a symmetric distribution around zero. A threshold of p-value=0.05 is used to denote significance which can only be achieved if there are a minimum of 6 comparisons per test.

#### Extraction Efficiency

In order to assess the efficiency of the minimally destructive extraction method, we first compare the proportion of endogenous molecules (i.e. molecules that align to the human reference genome, hg19) in samples produced using each extraction method and sequenced via shotgun (i.e. pre-capture) sequencing. While we observe a high degree of variability (Figure 2a; Table 1) between treatment types for each individual, there is a statistically significant difference in the proportion of endogenous molecules sequenced using the MDE and WTR methods (p-value=0.004), with an average of 35.8% and 18.8% endogenous molecules for each extraction method, respectively. These results support previous assertions that the outer cementum layer of the tooth root, which is targeted by the MDE method, contains a higher proportion of endogenous molecules than other portions of the tooth root (Damgaard et al. 2015). In contrast, we do not observe a significant difference in the proportion of endogenous molecules between methods MDE and P (p-value=1.000) (Supplementary Figure 2a), with an average of 36.4% endogenous observed when sampling from the petrous. These results are again consistent with claims that the petrous and tooth cementum both contain relatively high proportions of endogenous molecules (Damgaard et al. 2015; Hansen et al. 2017). While the high proportion of endogenous molecules obtained using the MDE method is promising, measuring the fraction of endogenous molecules in a sample does not tell us about the total amount of DNA obtained using each method.

**Figure 2.**
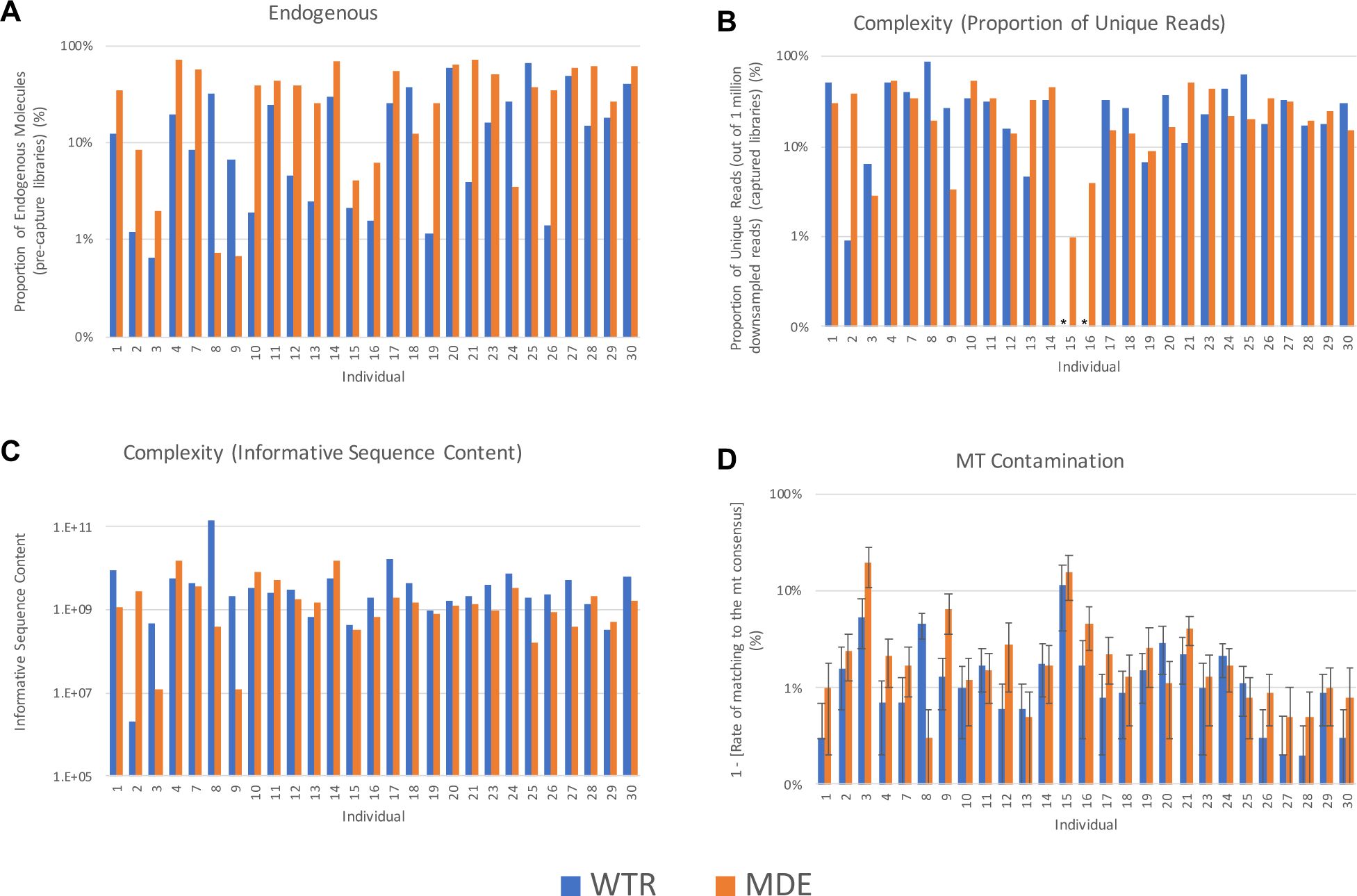
Sample Quality. A comparison of the quality of data produced by WTR (Whole Tooth Root) and MDE (Minimally Destructive Extraction) Methods in samples that passed quality filtering. (A) The proportion of endogenous molecules in data obtained via shotgun sequencing. (B) The complexity of each sample, as measured by the proportion of unique reads out of 1,000,000 reads sequenced. Asterisks indicate that the total number of unique reads sequenced was below 1,000,000 for the specified sample, therefore complexity estimates could not be generated. (C) The complexity of each sample, as measured by informative sequence content (D) The rate of contamination is compared by considering the rate of matching to mitochondrial consensus sequence. Error bars indicate the 95% confidence interval. Only samples that passed quality screening are shown. Plots showing comparisons with samples generated using Method P are shown in Supplementary Figure 2.

We therefore consider the overall complexity—the number of unique molecules contained within a single library—using two metrics. In the first metric, we consider the proportion of unique molecules sequenced in each sample, after down-sampling to 1,000,000 sequences that align to the 1.24 million SNPs targeted during capture (Figure 2B). This is a useful metric for comparison between samples, as it is not biased by differences in sequencing depth across samples. However, as this metric is calculated using sequence data for samples that underwent targeted enrichment capture, a process that may introduce bias into the data, we therefore also consider a second complexity metric, the informative sequence content (Glocke and Meyer 2017). This metric quantifies the relative proportion of molecules that were successfully amplified from each sample using quantitative PCR (qPCR) analysis. The results are calibrated using the proportion of endogenous molecules and average length of molecules measured in the shotgun sequencing data, reflecting the number of sequences in the DNA extracts that can be aligned to the human genome.

Neither complexity metric finds a statistically significant difference between complexity measured in samples prepared using Method MDE versus Method WTR (p-value=0.792 and 0.107, for the first and second complexity metrics, respectively), suggesting that using a minimally destructive extraction method does not result in loss of genetic data when sampling from teeth (Figure 2B, Table 1). While we find no statistically significant difference between samples prepared using Method MDE versus Method P using the first complexity metric (p-value=0.091), we do detect a significant difference using the second metric (p-value=0.043) (Supplementary Figure 2B-C). We note that the power of this analyses is limited due to the low number of comparisons we were able to make (N=7), therefore this comparison may warrant further study, particularly because previous studies have found that the rates of ancient DNA preservation in cementum versus petrous samples is dependent upon sample preservation (Hansen et al. 2017).

#### Contamination Rate

We were concerned that extracting ancient DNA directly from the outer layer of the tooth root might result in a higher rate of contamination in the sample, especially due to the increased potential for exposure of this region to contaminants during handling. Standard sampling protocols typically involve the physical removal of the outermost layer of bone or tooth prior to sampling, using a sanding disc or a sandblaster, while, in contrast, the minimally destructive extraction method specifically targets this outer layer following a superficial chemical (bleach) and brief (5-10 minute) ultraviolet decontamination. We therefore compare the relative contamination rates between sampling methods using a variety of metrics. First, we compare the rate of matching to the mitochondrial consensus sequence (Fu et al. 2013). A minimum threshold of 95% is typically applied during screening of ancient DNA for population genetic studies. We observe substantial variability in contamination rate between and within individuals for all treatment methods (Figure 2D, Table 1). While we detect a significant difference between mitochondrial match to consensus rates between the MDE and WTR methods (p-value=0.004), the average difference between these two methods is small (97.0% and 98.2%, respectively). Further, we observe no significant difference between the Methods MDE and P (p=0.310) (Supplementary Figure 2D).

Next, we estimate the autosomal rate of contamination, using the tool ContamLD (Nakatsuka et al. 2020), which measures the breakdown of linkage disequilibrium in a sequenced individual, a process which is accelerated by increased contamination. We again estimate relatively low rates of contamination across all samples, and find no significant difference in contamination rates between Methods MDE and WTR (p-value=0.490) or between Methods MDE and P (p-value=0.893).

We also estimate contamination rates in the individuals who are identified as genetically male using ANGSD (Korneliussen et al. 2014). We obtain low estimates of contamination (≤2.5%) across all male samples (Table 1). Comparing the X-chromosome contamination estimates for the 6 genetically male individuals for whom there was enough data to produce estimates for both treatment types, we do not detect a significant difference between the MDE and WTR Methods (p-value=0.293). Taken together these three estimates of contamination suggest that, in practice, the UV and bleach decontamination protocol used for the MDE Method performs similarly to the physical surface removal decontamination steps implemented in the destructive protocols, and is sufficient to produce ancient DNA data of analyzable quality.

We considered the read length distribution and frequency of C-to-T damage in the terminal bases of reads that aligned to the human genome (hg19) that were obtained via shotgun sequencing (i.e. pre-capture). Authentic ancient DNA is thought to consist of characteristically short fragments, with very few reads longer than 100 base pairs (Sawyer et al. 2012; Dabney et al. 2013b; Glocke and Meyer 2017), therefore the read length distribution is used as a general metric to assess ancient DNA authenticity. We find that all samples appear to have read length profiles characteristic for authentic ancient DNA (Supplementary Figure 3) and we do not observe a significant difference in median length of reads obtained using Method MDE and Method WTR (p-value=0.375). A weakly significant difference is observed between reads obtained using Method MDE and P (p-value=0.034) (Table 1), suggesting that there may be systematic differences between DNA preservation in petrous and tooth samples.

Endogenous ancient DNA samples are also thought to exhibit a high rate of C-to-T damage, particularly in the terminal bases. Using a partial or USER UDG treatment for double stranded and single stranded libraries, respectively (Rohland et al. 2015; Gansauge et al. in Prep), we removed this damage in the interior of each molecule, while retaining it in the terminal bases. Therefore, we are able to use the frequency of these errors to assess ancient DNA authenticity. For samples processed using Method MDE and WTR (p-value=0.249) we observe no significant difference in the frequencies of C-to-T damage in terminal bases at the 5’ end of molecules that aligned to the human genome (hg19), obtained via shotgun sequencing. However, the distribution of damage rates in samples processed using Method P are significantly different to Method MDE (p-value=0.028), with higher rates of damage observed in libraries produced using Method P in most (8/9) cases, again suggesting that there may be systematic differences between DNA preservation in petrous and tooth samples (Table 1, Supplementary Figure 4 & 5).

Finally, we were concerned that the use of parafilm to cover portions of the tooth roots that we did not want expose to the extraction buffer could serve as a possible source of contamination. We therefore created a parafilm extraction control, in which a small strip of parafilm (comparable in size to that used for covering the tooth roots), was added to a tube of extraction buffer and underwent sample processing along with the MDE samples and regular extraction blanks. We observe very few reads associated with this parafilm blank (Supplementary Table 1), suggesting that the use of parafilm does not serve as a significant source of contamination in the MDE Method.

## DISCUSSION

This minimally destructive sampling protocol enables extraction of ancient DNA from the cementum portion of tooth roots that is of similar quality to ancient DNA obtained from teeth using traditional, destructive sampling methods that rely on powder produced through drilling or cutting and powdering. This is true with regards to both the amount of DNA that it is possible to obtain and the levels of contamination detected in the samples. In contrast, our results suggest that DNA sampled from the petrous bone exhibits more complexity than DNA sampled from the tooth cementum, indicating that there is still justification for choosing to sample from petrous bones over teeth when trying to maximize the chances of successfully sequencing ancient DNA, particularly in cases where sample preservation is poor—a circumstance in which ancient DNA sampled from petrous has previously been found to be of higher quality than in cementum (Hansen et al. 2017). However, the physical damage to the sampled tooth is substantially reduced and the morphological integrity of the sampled tooth is retained when using this minimally destructive sampling protocol, making this an optimal sampling method of teeth in cases where sample preservation is of the highest priority.

One of the major concerns surrounding an extraction protocol that targets the outer surface of an ancient sample is the potential for an increase in contamination, as this outer surface may come in direct contact with various contaminants, particularly during handling. Since the majority of samples selected for ancient DNA analysis have been excavated and manipulated without any consideration for potential future genetic studies, this is of particular concern. While destructive methods physically remove the outermost layer of bones and teeth to reduce contamination, we instead applied a bleach and UV decontamination procedure to the tooth before processing. We detected little difference in contamination rates between samples processed using this minimally destructive decontamination and sampling method and those processed using standard destructive methods. Further, these results suggest that decontamination procedures that involve wiping a sample with bleach do not significantly reduce DNA yields, as opposed to previously proposed decontamination methods involving the soaking of the sample for an extended period of time (e.g. Higgins et al. 2013). By targeting the outer cementum tooth surface directly, this method maximizes the proportion of cementum matrix which is being digested and minimizes the amount of dentine sampled when compared to other cementum-targeting methods (Damgaard et al. 2015), which sample a significant proportion of the inner dentine layer in addition to the cementum. Furthermore, we find that parafilm can be used to protect portions of the tooth that users do not wish to sample (i.e. the tooth crown) from exposure to extraction buffer, without increasing contamination rates.

While these results show that this minimally destructive approach is a promising alternative to destructive sampling methods that are traditionally applied to ancient teeth, we stress that further research is needed to determine whether it is recommended to opt for this sampling method in all circumstances. Particularly, we note that the majority of teeth chosen for this analysis were of moderate to excellent preservation status. The two most poorly preserved individuals included in this study contained too little DNA to allow for comparisons to be made between Methods MDE and WTR, and the tooth roots processed via Method MDE sustained damage during processing. Further study of the utility of this method on less well-preserved teeth is therefore of great interest.

As the impact on dental morphology is minimal, this approach enables the preservation of samples for future analyses. Previous studies have shown that exposure to the chemicals used for ancient DNA extraction (mainly EDTA and proteinase K) do not affect a specimen’s suitability for subsequent biochemical analyses, such as radiocarbon (AMS C14) dating (Korlević et al. 2018). Therefore, teeth processed using this minimally destructive protocol would remain suitable for future biochemical analyses.

This minimally destructive extraction method drastically reduces the amount of physical destruction caused by ancient DNA extraction, creating no holes or cuts in the sampled tooth or bone, while also shortening the overall length of the extraction protocol, without meaningfully increasing the amount of contamination. This method makes it possible to extract ancient DNA from individuals that would otherwise be unavailable for ancient DNA study due to the destructive nature of traditional sampling methods.

## METHODS

All ancient DNA analyses were performed in dedicated clean rooms at the University of Vienna and Harvard Medical School. For individuals 1-10, skeletal sampling, preparation and DNA extraction were performed at the University of Vienna. Library preparation, targeted enrichment capture, and sequencing was performed at Harvard Medical School. For individuals 11-30, skeletal sampling was performed at the University of Vienna, while all other processing was performed at Harvard Medical School.

### Sampling

We selected skeletal elements from 30 ancient individuals of varying age, geographic origin, and degree of preservation for analysis (Table 1). From each individual, we selected a single multi-rooted tooth for sampling. For the first 10 individuals, we also selected a temporal bone for sampling. We UV irradiated each tooth in a cross-linker for 5 to 10 minutes on each side, in order to remove as much surface contamination as possible. We then cut off the roots of each tooth using a diamond cutting disc and a hand-held Dremel drill, treating each root separately in all subsequent analyses. From each individual, we randomly selected one tooth root (“Method MDE”) for minimally destructive extraction. These tooth roots were subject to additional surface cleaning by wiping the teeth clean with a 2% bleach solution and rinsing with 95% ethanol, followed by UV-irradiation for 5 to 10 minutes on each side. We prepared the second set of tooth roots (“Method WTR”) by removing the extreme outer surface of each tooth root using a sanding disc and drill, and milling the root in a Retsch MM400 mixer mill for a total of 60 seconds with a 10 seconds break after 30 seconds to produce a powder. Additionally, we obtained approximately 50mg of bone powder from the petrous portion of each of the 10 selected temporal bones, using standard methods (“Method P”) (Pinhasi et al. 2019).

### DNA Extraction

We prepared selected tooth roots (Method MDE) for minimally destructive extraction by recording the initial weight of the tooth root, then isolating the targeted portion of the tooth root using parafilm (Supplementary Figure 6; see Supplementary Information 1 for a step-by-step description of the minimally destructive extraction method). We targeted the lower portion of the tooth root, where cellular cementum is concentrated. All other surfaces were wrapped in UV-decontaminated parafilm in order to prevent significant contact with the extraction buffer. The tooth roots were placed in 750 µL −1 mL of extraction buffer (0.45 M EDTA, 0.25 mg/mL Proteinase K, pH 8.0; defined in Rohland and Hofreiter (2007) with the exposed portion pointing down, and incubated for 2.5 hours at 37°C, shaking gently. Following incubation, the roots were removed from the extraction buffer, which was then processed according to standard ancient DNA extraction procedures. Samples from individuals 1-10 underwent manual ancient DNA extraction, as described in Dabney et al. (2013a), with modifications. The MinElute columns were replaced with a preassembled spin column device (Roche, as described in Korlević et al. (2015)). We washed lysates with 650 µL of PE buffer (Qiagen) and spun at 6000 rpm for 1 minute. Following dry spin, we isolated the DNA by placing the spin column in a fresh 1.5 mL collection tube, and 25 µL TET buffer was pipetted onto the column’s silica membrane, which was incubated at room temperature for 10 minutes, and then spun at maximum speed for 30 seconds. We repeated this step, producing a total of 50 uL of DNA extract. Samples from individuals 11-30 underwent robotic extraction following incubation, using the robotic protocol described in Rohland et al. (2018), using buffer D.

For samples processed using Methods WTR and P, sampled bone powders were incubated overnight (~18 hours) in extraction buffer at 37°C, with gentle shaking. For samples from individuals 1-10, up to 50mg bone powder was incubated in 1mL extraction buffer, which then underwent manual extraction, as described above. For samples from individuals 11-30, ~37 mg of bone powder was incubated in 750 μL extraction buffer, and then underwent robotic extraction, as described above.

Negative controls were prepared alongside ancient DNA extracts for all extraction batches. In each case, extraction buffer was added to an empty tube prior to incubation, and the negative control was treated identically to all other samples during subsequent processing. Additionally, we generated one parafilm extraction control, by incubating a piece of UV-decontaminated parafilm in extraction buffer overnight in order to determine whether the parafilm coverings used to protect the ends of the tooth roots might be a potential source of contamination.

Following incubation in the extraction buffer, the roots were rinsed with 95% ethanol in order to remove any remaining extraction buffer and air dried at room temperature for 24 hours. The samples were then re-weighed to assess the total amount of dental material digested.

### Library Preparation, Enrichment, and Sequencing

We prepared double-stranded (samples 1-10) or single-stranded (samples 11-30) libraries from 10 µL of each extract using UDG-treatment methods, as described in Rohland et al. (2015) and Gansauge et al. (in Prep), respectively. These methods remove ancient DNA damage at the interior of each DNA sequence, while preserving characteristic ancient DNA damage at the terminal ends of the molecules, to be used for ancient DNA authentication during bioinformatic processing. We enriched libraries for human DNA via targeted enrichment at 1.24 million SNP sites that are informative for population genetic analyses (Fu et al. 2015; Haak et al. 2015; Mathieson et al. 2015). Following enrichment, libraries were sequenced on an Illumina NextSeq500 machine, with 2×76 or 2×101 cycles, with an additional 2×7 or 2×8 cycles used for identification of indices, for double-stranded and single-stranded libraries, respectively.

### Bioinformatic Processing

We trimmed molecular adapters and barcodes from sequenced reads, and the merged paired end reads, requiring an overlap of 15 base pairs (allowing up to three mismatches of low base quality (<20) or one mismatch of high base quality (≥20)) using custom software (https://github.com/DReichLab/ADNA-Tools). We then aligned the merged sequences to both the mitochondrial RSRS genome (Behar et al. 2012) and the hg19 human reference sequence using *samse* in bwa (v0.6.1) (Li and Durbin 2009). We identified duplicate reads, defined as having the same start and end position and orientation, and a shared DNA barcode (unique quadruple barcode combinations are inserted during library preparation), and retained only the copy with the highest quality sequence.

We assessed ancient DNA authenticity using several metrics. We used the tool ContamMix (Fu et al. 2014) to determine the rate of matching between mitochondrial reads and the consensus sequence. The tool ContamLD was used to estimate the rate of contamination in the autosomes, based on the degree of breakdown of linkage disequilibrium observed in each library relative to a panel of representative individuals from the 1000 Genomes project (Nakatsuka et al. 2020). We determined the amount of contamination in the X-chromosome for male individuals using the tool ANGSD (Korneliussen et al. 2014). Finally, we estimated the rate of C-to-T substitution at the terminal ends of molecules for each sample (Jónsson et al. 2013) and the lengths of sequenced molecules were considered as metrics of DNA authenticity for each sample.

We assessed the quality of ancient DNA observed by measuring the percent of endogenous (unique reads that align to the human genome), coverage (average number of reads aligning to each of the 1.24 million targeted SNP sites), and overall complexity of the sample—assessed by determining the proportion of unique reads sequenced, after randomly down-sampling to 1,000,000 on-target reads, or by measuring the informative sequence content (Glocke and Meyer 2017), in order to minimize bias caused by differences in sequencing depth.

## ACKNOWLEDGEMENTS

We thank Iñigo Olalde and Nathan Nakatsuka for contributions to the bioinformatic analyses. E.H. was supported by a graduate student fellowship from the Max Planck-Harvard Research Center for the Archaeoscience of the Ancient Mediterranean (MHAAM). D.R. is an Investigator of the Howard Hughes Medical Institute and this work was also supported by John Templeton Foundation grant 61220. Tamás Hajdu, Tamás Szeniczey, János Dani and Krisztián Kiss were supported by grant from the Hungarian Research, Development and Innovation Office, project number: FK128013. Alexandra Anders was supported by a grant from the Hungarian National Research, Development and Innovation Fund (Grant K124326).

