## Supplementary Information for "A Minimally Destructive Protocol for DNA Extraction from Ancient Teeth"

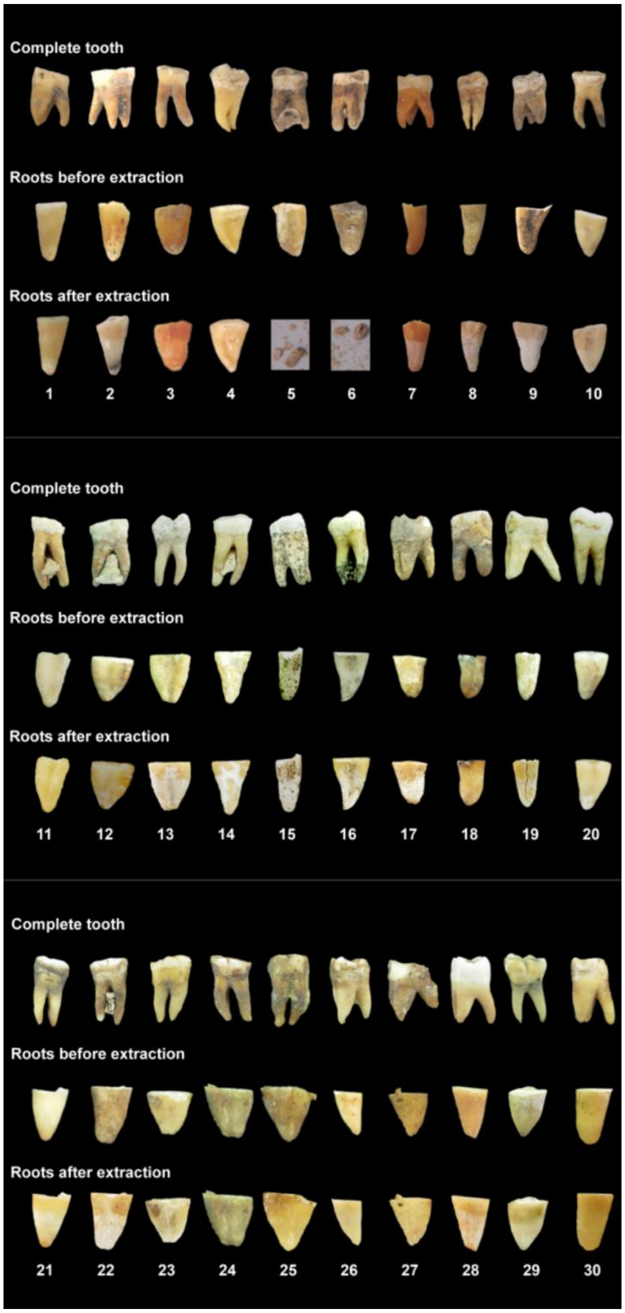

### Supplementary Figure 1. Tooth root before and after minimally destructive extraction.

The complete tooth is shown prior to processing (top row). Tooth roots are shown immediately prior to extraction (middle row) and 24 hours after extraction (bottom row).

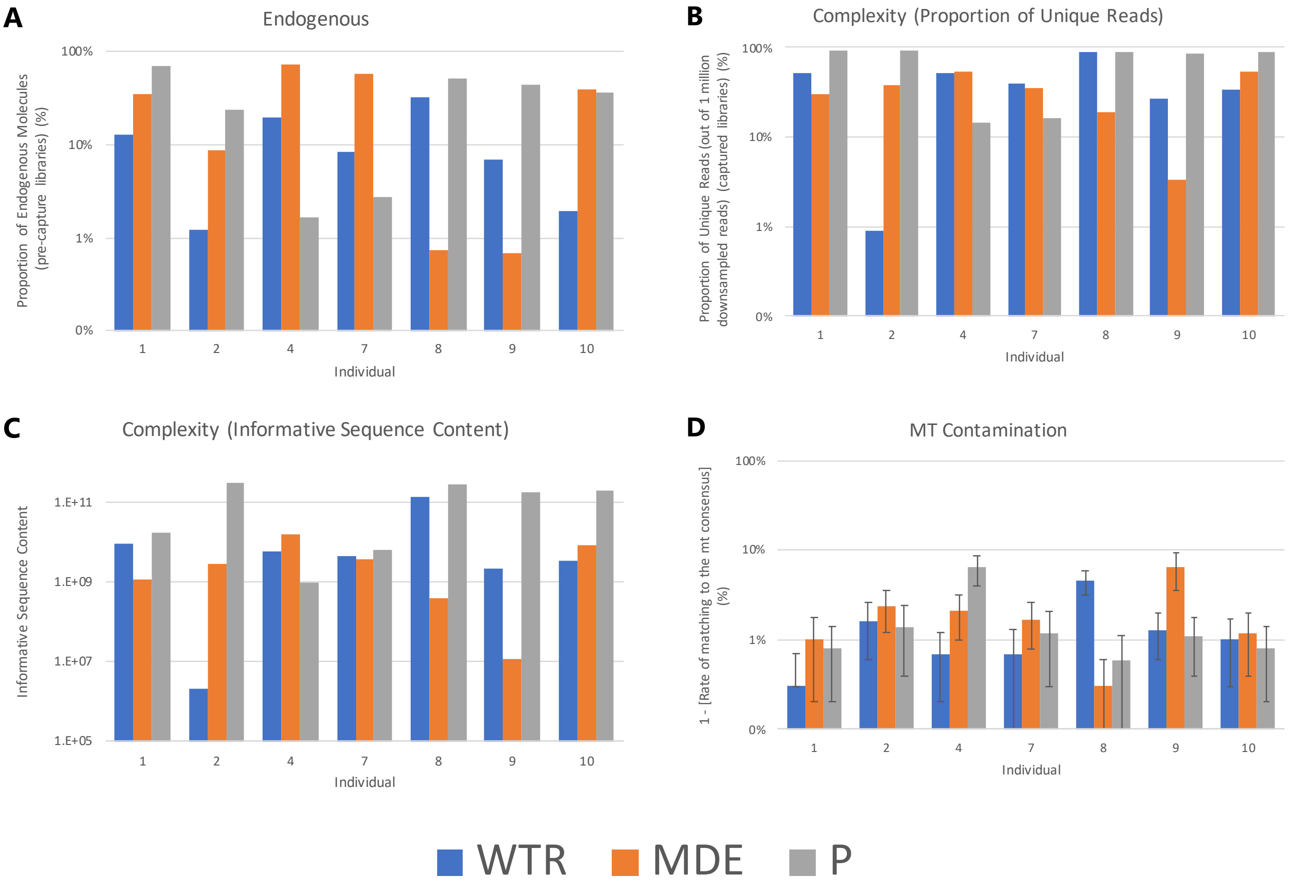

### Supplementary Figure 2. Sample quality with petrous comparisons

A comparison of the quality of data produced by Method MDE (Minimally Destructive Extraction), Method WTR (Whole Tooth Root) and Method P (Petrous) in samples that passed quality filtering and that had data available for all three treatment types. (A) The proportion of endogenous molecules in data obtained via shotgun sequencing is compared. (B) The complexity of each sample, as measured by the proportion of unique reads out of 1,000,000 reads sequenced. (C) The complexity of each sample, as measured by informative sequence content (D) The rate of contamination is compared by considering the rate of matching to mitochondrial consensus sequence. Error bars indicate the 95% confidence interval.

### Double Stranded Libraries

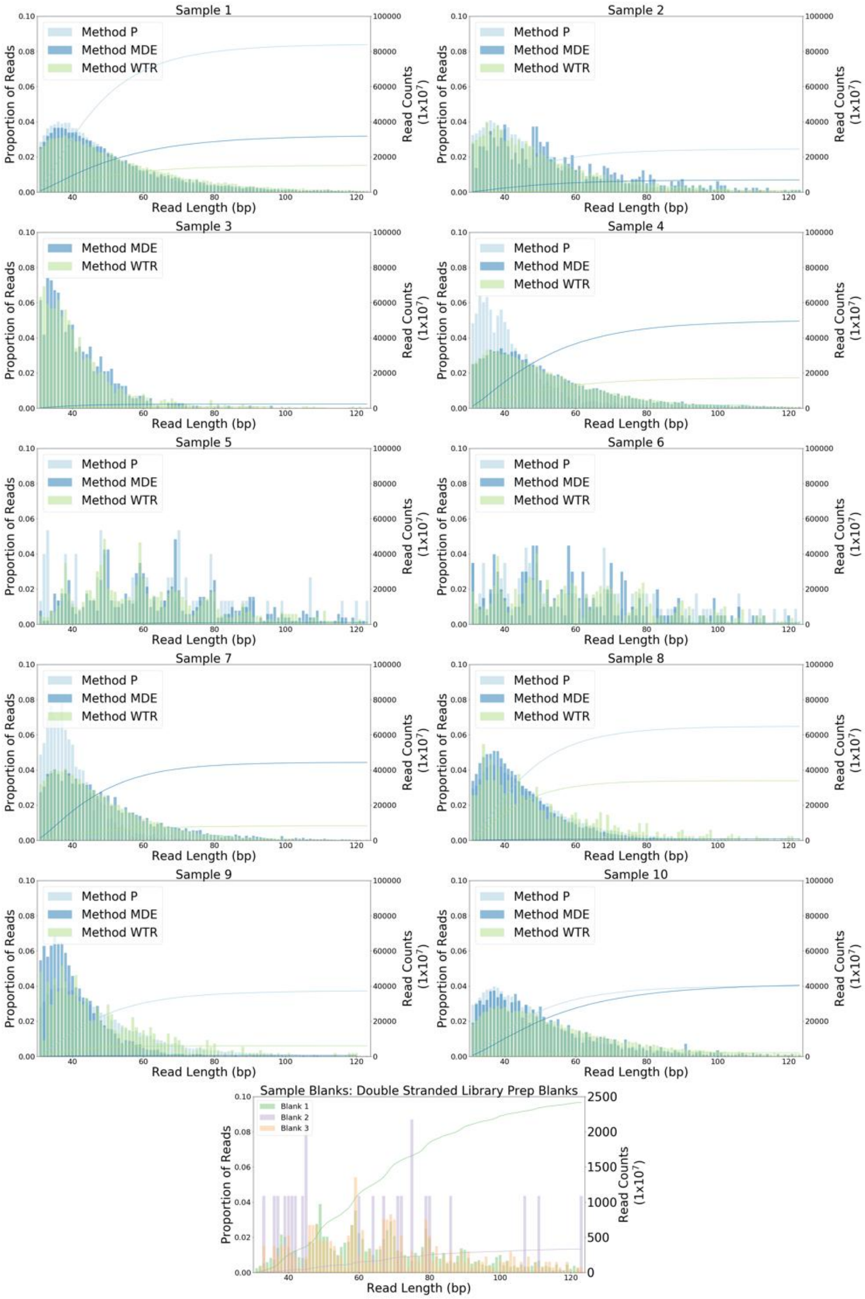

### Single Stranded Libraries

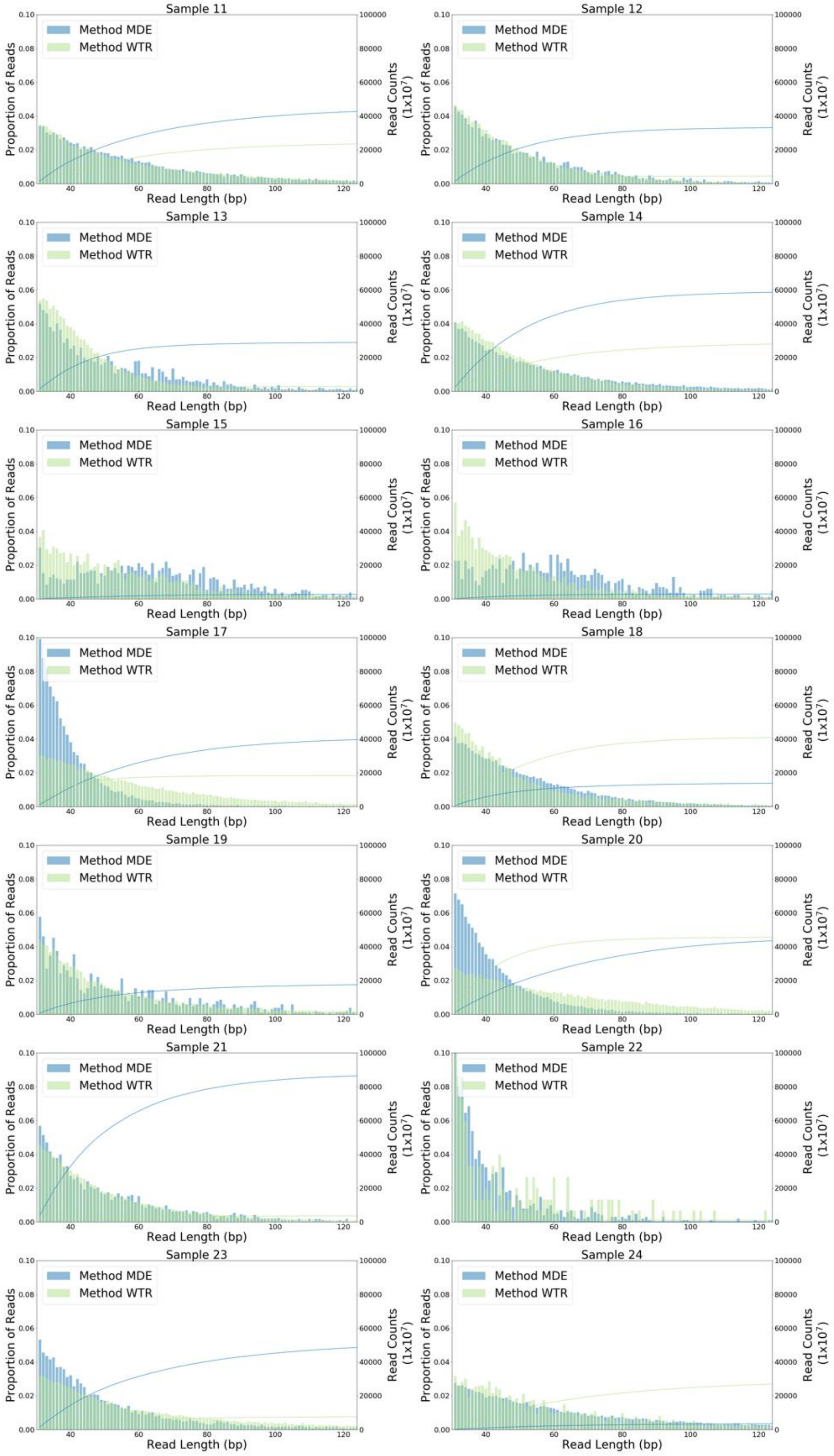

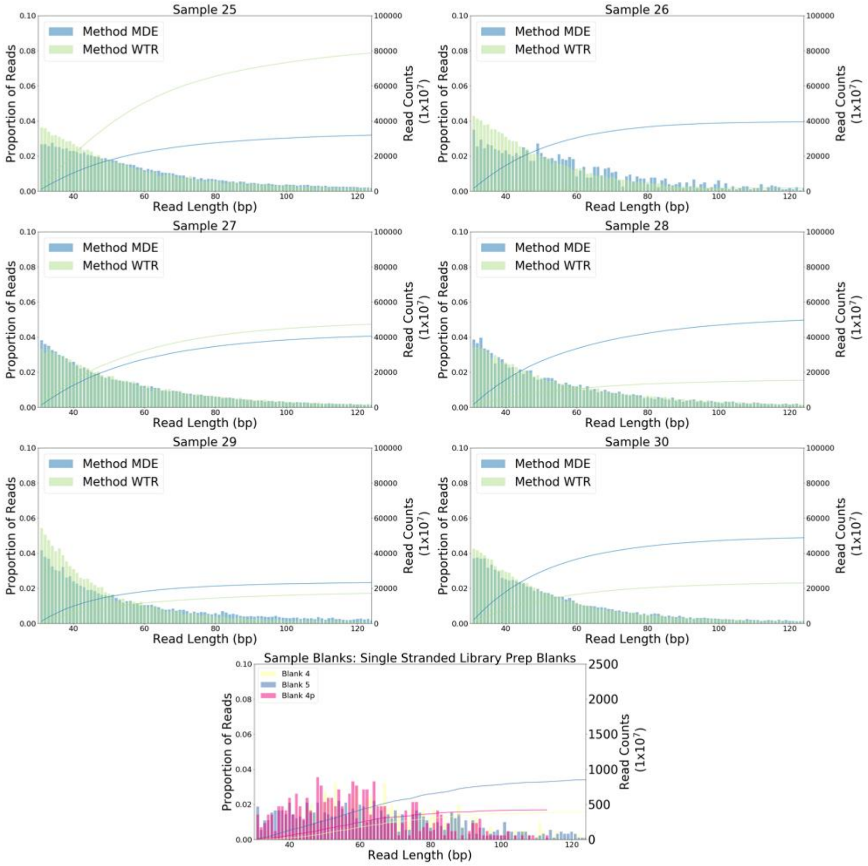

### Supplementary Figure 3. Read length distribution.

A histogram showing the distribution of molecule lengths for reads sequenced pre-capture (i.e. shotgun sequencing) that align to the human genome (hg19), for extraction methods MDE, WTR and P. The cumulative distribution of reads is also shown, with the y-axis on the right showing the total number of reads.

### Double Stranded Libraries

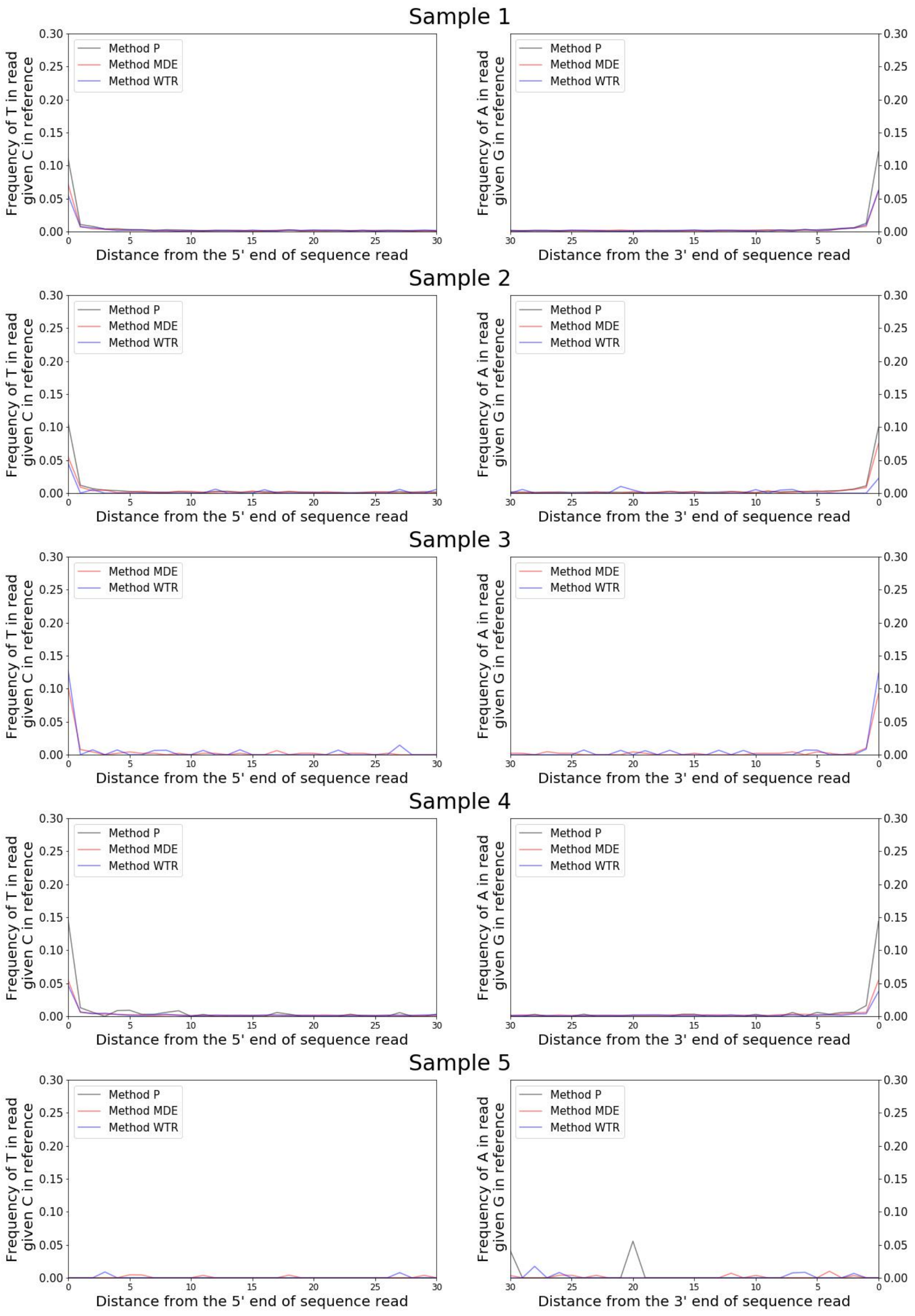

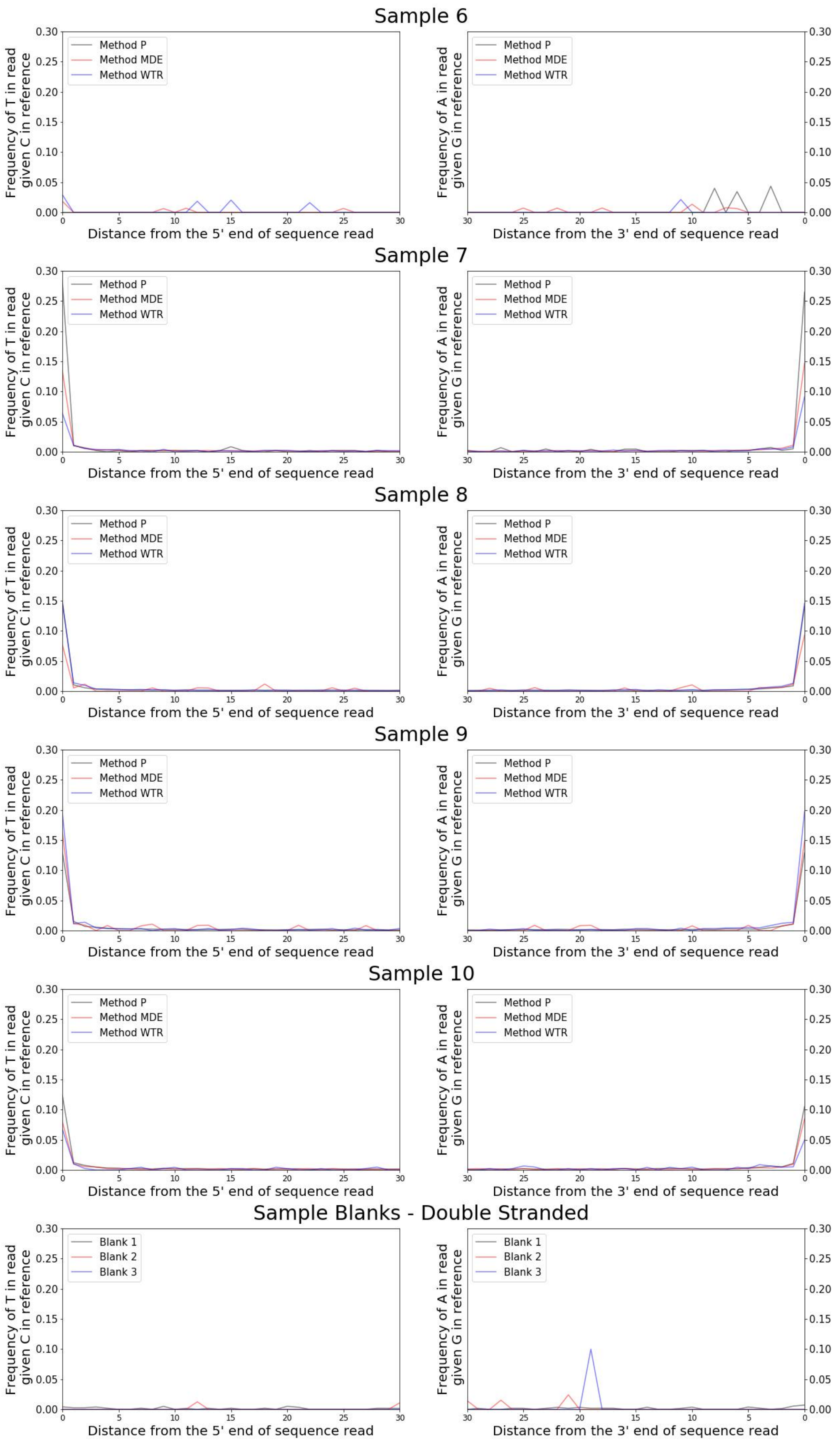
Sample 6
Sample 7
Sample 8
Sample 9
Sample 10
Sample Blanks - Double Stranded
Frequency of T in read given C in reference
Frequency of A in read given G in reference
Distance from the 5' end of sequence read
Distance from the 3' end of sequence read
Method P
Method MDE
Method WTR
Blank 1
Blank 2
Blank 3

### Single Stranded Libraries

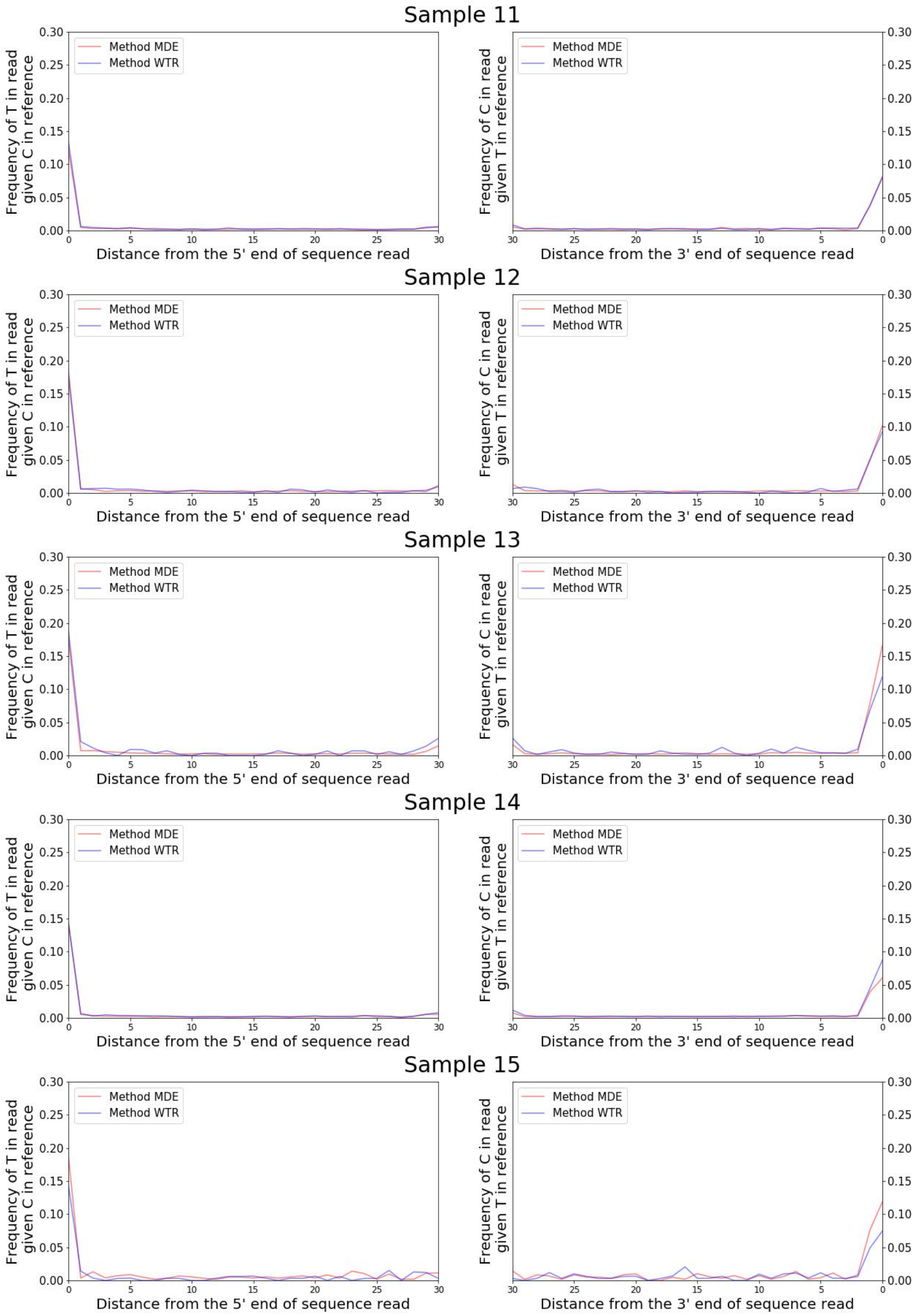

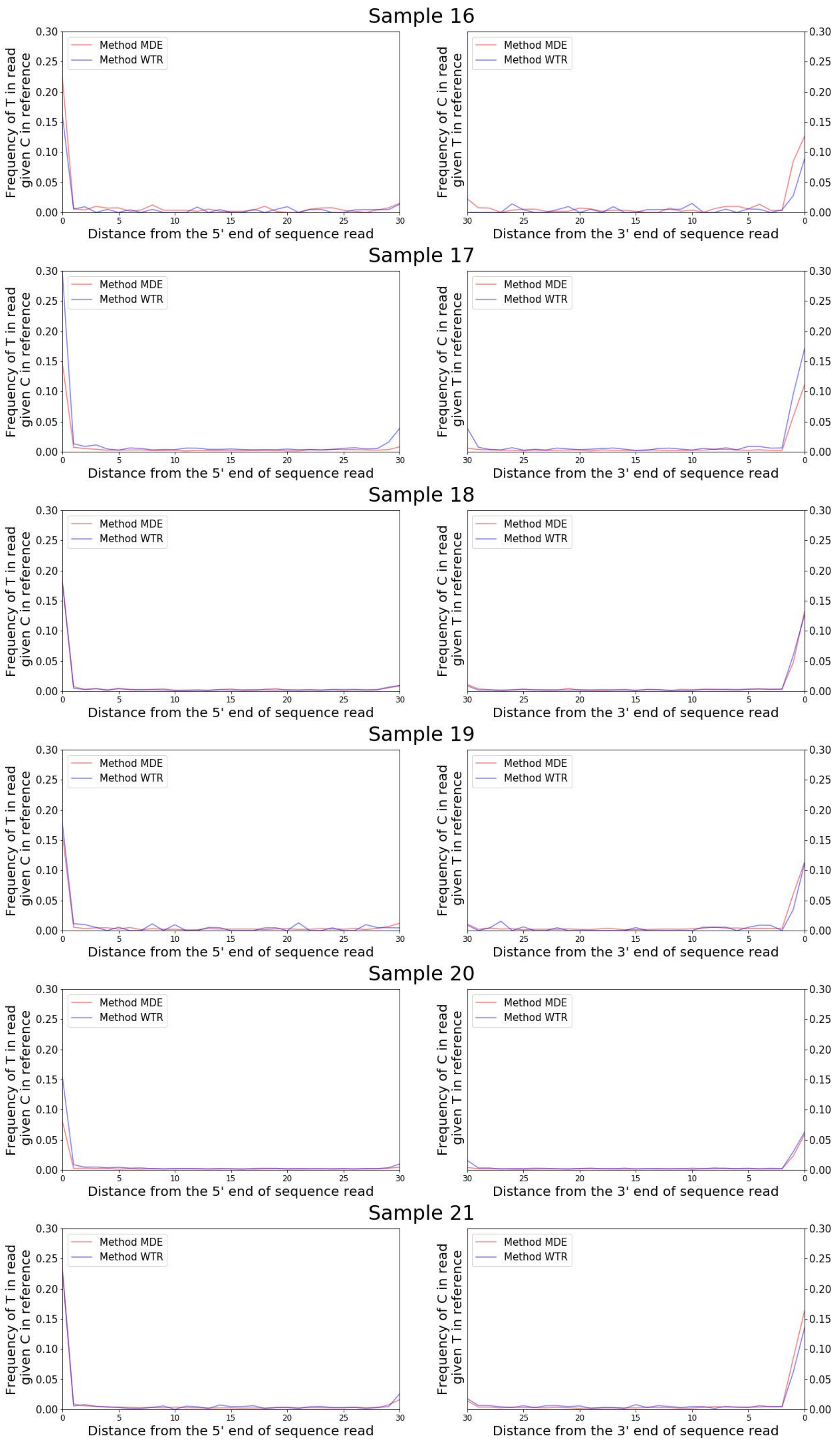
Sample 16
Sample 17
Sample 18
Sample 19
Sample 20
Sample 21
Method MDE
Method WTR
Frequency of T in read given C in reference
Frequency of C in read given T in reference
Distance from the 5' end of sequence read
Distance from the 3' end of sequence read

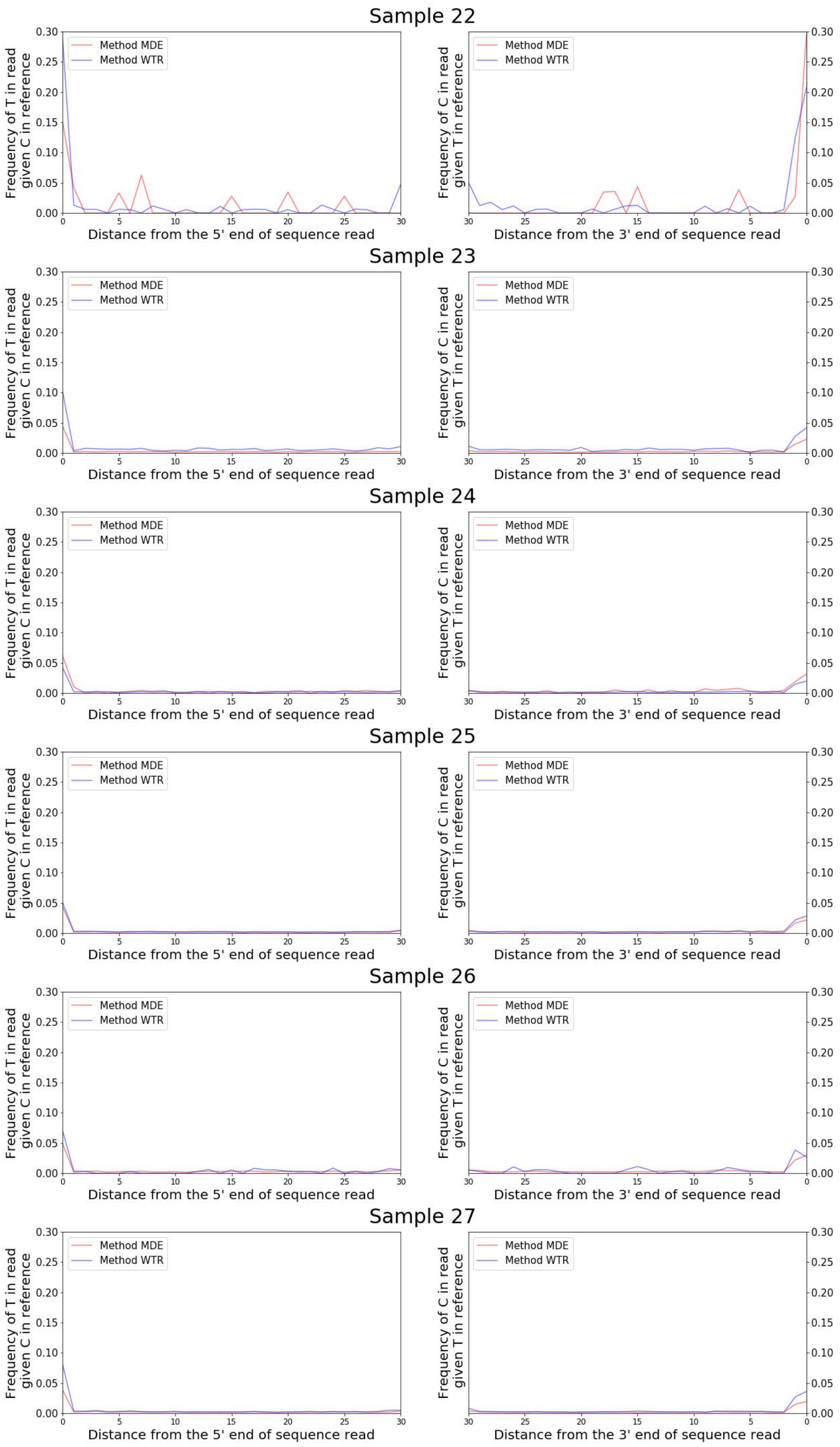
Sample 22
Method MDE
Method WTR
Frequency of T in read given C in reference
Distance from the 5' end of sequence read
Frequency of C in read given T in reference
Distance from the 3' end of sequence read
Sample 23
Method MDE
Method WTR
Frequency of T in read given C in reference
Distance from the 5' end of sequence read
Frequency of C in read given T in reference
Distance from the 3' end of sequence read
Sample 24
Method MDE
Method WTR
Frequency of T in read given C in reference
Distance from the 5' end of sequence read
Frequency of C in read given T in reference
Distance from the 3' end of sequence read
Sample 25
Method MDE
Method WTR
Frequency of T in read given C in reference
Distance from the 5' end of sequence read
Frequency of C in read given T in reference
Distance from the 3' end of sequence read
Sample 26
Method MDE
Method WTR
Frequency of T in read given C in reference
Distance from the 5' end of sequence read
Frequency of C in read given T in reference
Distance from the 3' end of sequence read
Sample 27
Method MDE
Method WTR
Frequency of T in read given C in reference
Distance from the 5' end of sequence read
Frequency of C in read given T in reference
Distance from the 3' end of sequence read

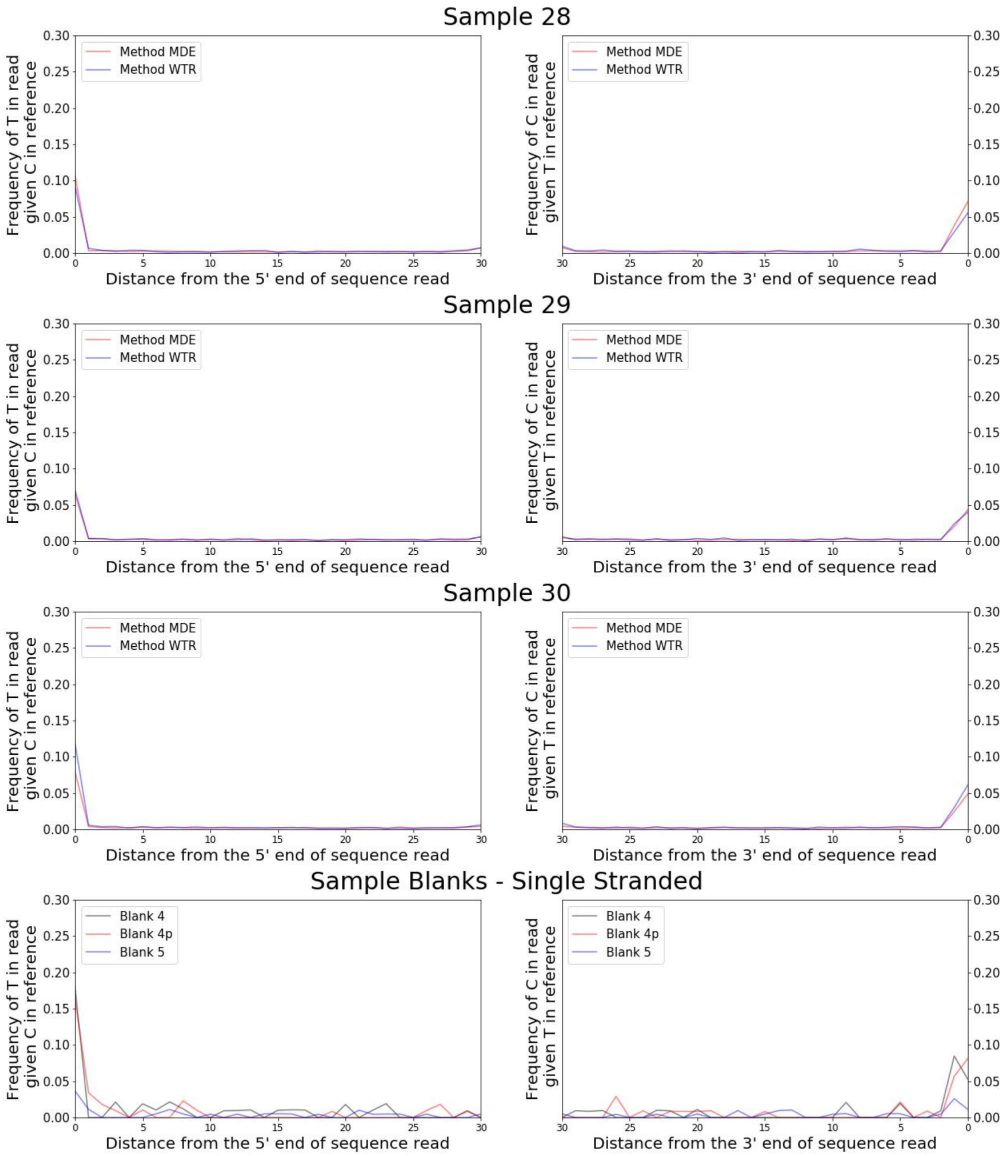

### Supplementary Figure 4. C-to-T damage profiles

Damage profiles calculated and plotted for each individual using mapDamage2.0 (Jónsson et al, 2013). For each individual, we plot the frequency of misincorporations at the 30 terminal base pairs of the molecule for all libraries for pre-capture (i.e. shotgun sequencing) reads that align to the human genome (hg19). For double stranded libraries (samples 1-10), we report the rate of C-to-T misincorporations at the 5’ end and G-to-A misincorporations at the 3’ end of reads (plotted on the left and right respectively). For single stranded libraries (samples 11-30), we report the rate of C-to-T misincorporations at both the 5’ and 3’ ends of reads (plotted on the left and right respectively).

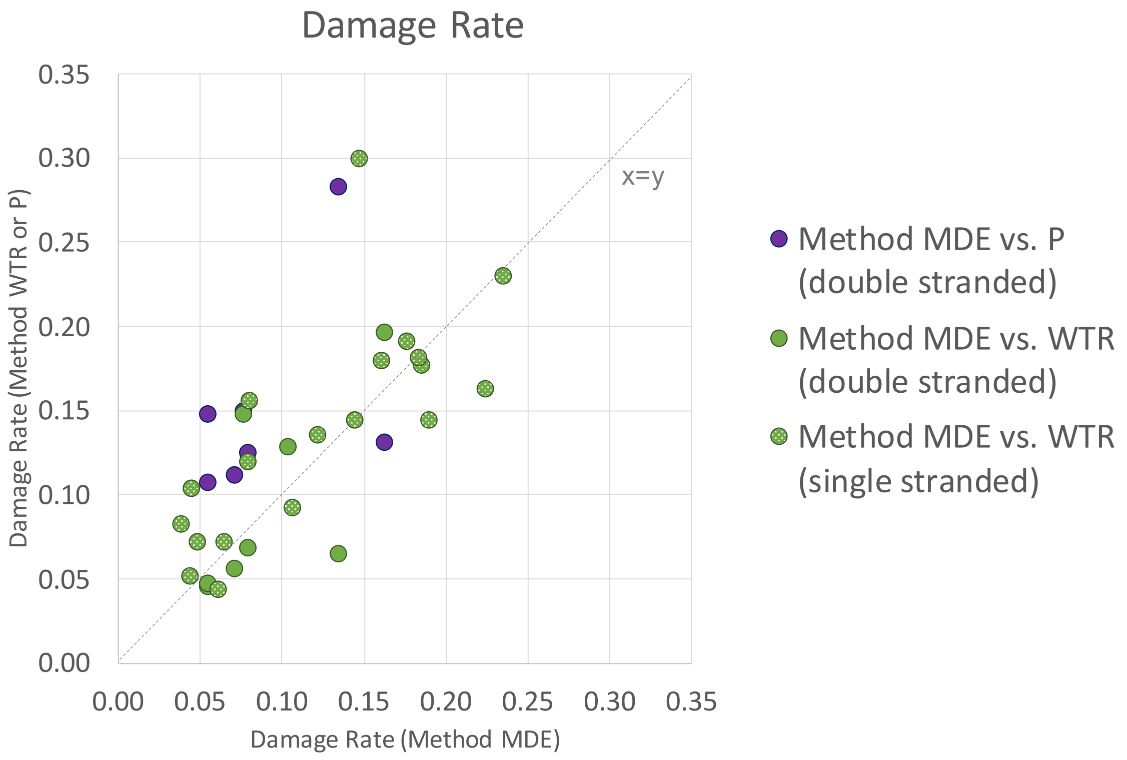

#### Supplementary Figure 5. C-to-T Damage Rate Comparison

A scatterplot comparing the 5' C-to-T damage rates at the terminal ends of each molecule for libraries prepared using Method MDE vs Method WTR or P, that aligned to the human genome (hg19) following pre-capture (i.e. shotgun) sequencing. Double stranded libraries are represented with solid colored markers, while single stranded libraries are represented with patterned markers. The line y=x is shown in grey.

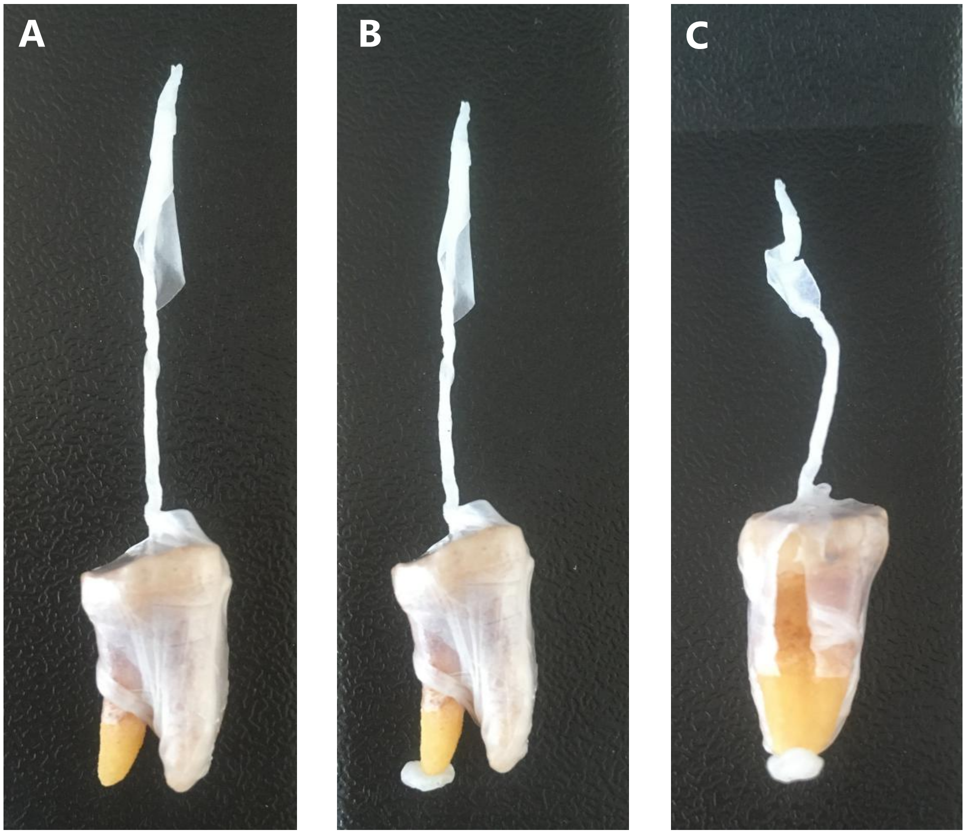

#### Supplementary Figure 6. Example tooth wrapped in parafilm

[A] Picture of a tooth that has been wrapped in parafilm leaving only the lower part of one of the roots exposed. Note the tail of parafilm present to allow for easier handling of the tooth. [B] A small piece of parafilm has been pressed to the bottom of the tooth root to cover any visible holes that may be present. [C] The parafilm wrapping has been sliced using a metal blade, demonstrating one method for removing the parafilm wrapping following extraction.

### SUPPLEMENTARY INFORMATION 1
### Detailed Minimally Destructive Extraction Protocol – manual processing

#### Required Reagents and consumables:

| Reagent/Consumable | Supplier (Austria) | Cat. No. (Austria) |
|---|---|---|
| Water | Carl Roth | 3255.1 |
| EDTA (0.5M) | VWR | E177-500ML |
| Guanidine hydrochloride | Glentham Life Sciences | GE1914 |
| Isopropanol | Fisher | 10284200 |
| Sodium Acetate (3M, pH 5.2) | Alfa Aesar | J61928.AK |
| Tween-20 | Sigma | P9416-50ML |
| Tris-HCl (1M, pH 8.0) | Fisher | BP1758-500 |
| PE Wash Buffer | Qiagen | 19065 |
| Parafilm | VWR | 291-1214 |
| 5mL microcentrifuge tubes | Eppendorf | 0030119460 |
| 50 mL conical tubes | VWR | 525-0610 |
| High Pure Viral Nucleic Acid Large Volume Kit (incl. Proteinase K) | Roche | 05114403001 |
| 1.5 mL DNA LoBind microcentrifuge tubes | Eppendorf | 0030108051 |

### Required Buffers:

#### Extraction Buffer:

| | Final Concentration | 1mL | 50mL |
|---|---|---|---|
| $H_20$ | .. | 83.3 µL | 45 |
| EDTA (0.5M) | 0.45 M | 900 µL | 4.165 mL |
| Proteinase K* | 0.25 mg/mL | 16.7 µL | 835 µL |

* Prior to adding proteinase K, UV-irradiate the solution in for 30 minutes

#### Binding Buffer*:

| | Final Concentration | 50mL |
|---|---|---|
| Guanidine hydrochloride | 5 M | 23.9 g |
| Isopropanol | 40% | 20 mL |
| Sodium Acetate (3M) | 90 mM | 1.5 mL |
| Tween-20 (10%) | 0.05% | 250 µL |
| $H_20$ | .. | Fill to 50 mL |

* Store for no longer than 1 month. UV-irradiate the solution 30 minutes before use.

#### TET Buffer:

| | Final Concentration | 50mL |
|---|---|---|
| Tris-HCl (1 M, pH 8.0) | 10 mM | 500 µL |
| EDTA (0.5 M) | 1 mM | 100 µL |
| Tween-20 (10%) | 0.05 % | 250 µL |
| $H_20$ | .. | 49.15 mL |

**Procedure:**

1. **Prepare Reagents and Materials:** Prepare all required buffers. Cut parafilm to the desired size (we recommend approximately 1.5 cm x 3 cm rectangle). UV irradiate all UV-insensitive reagents and materials for a minimum of 30 minutes prior to beginning extraction. NOTE: Proteinase K is UV-sensitive and should be added to the extraction buffer solution after UV irradiation.
2. **Physically Clean Tooth**: Wipe tooth gently but thoroughly with a solution of 2% bleach using a disposable paper wipe to remove surface contamination. Next rinse or wipe tooth with 95% ethanol to remove bleach. NOTE: If the tooth has been treated with varnish or glue, we recommend trying to identify the substance used on the tooth to determine which solvent would be the most suitable, possible options include 99.5% acetone, 95% ethanol and purified water. A cleaning step involving either water or acetone should be followed by 95% ethanol rinse before performing the bleach cleaning step.
3. **UV Irradiation:** UV irradiate tooth surface by placing it in a cross-linker or similar device for a minimum of 5-10 minutes at 254 nm. In order to be certain that the entire surface has been irradiated, turn the tooth after 5-10 minutes and continue with UV irradiation at 254 nm. After this process, ensure that the tooth is completely dry before proceeding to the next step.
4. **Prepare Tooth Surface:** Isolate the extraction surface (we recommend targeting the lower third of the tooth root) for extraction using parafilm. Tightly wrap film around crown of the tooth and upper root surface. In cases where a hole or crack is visible in the tooth root, also cover this with a small piece of parafilm NOTE 1: We recommend leaving a small "tail" of parafilm above the crown to make it easier to remove the tooth from the tube following incubation (Supplementary Figure 6). NOTE 2: We caution users not to be too overzealous when wrapping teeth in parafilm, particularly when sampling from very poorly preserved samples. The more parafilm is used, the more difficult it is to remove following extraction. This may result in sample breakage if too much pressure is applied to a very poorly preserved sample.

5. **Incubate Tooth in Extraction Buffer:** Place the prepared tooth in a 5mL tube with the exposed part of the root pointing down. Add 1 mL of extraction buffer. Digest for 2.5 hours at 37ºC, with gentle horizontal rotation, to allow extraction buffer to circulate around tooth root. NOTE 1: Ideal incubation time may vary depending on sample quality. An indicator of when an ideal incubation time has been reached may be when a difference in the surface level of the digested versus undigested material becomes visually apparent. NOTE 2: Be sure to use enough extraction buffer to completely cover the portion of the tooth that you are interested in sampling from. If additional extraction buffer is used, it is also necessary to proportionally increase the amount of binding buffer added to the sample in the next step.
6. **Prepare Binding Buffer:** Add 13mL of binding buffer to a 50mL conical tube.
7. **Remove Tooth from Extraction Buffer (Lysate):** Using a 100-1000 µL pipette tip, remove as much of the lysate from the 5mL tube as possible and transfer to the 50 mL conical tube containing binding buffer prepared in the previous step. Next, remove the tooth from the 5mL tube, and set it aside. If any lysate remains in the 5 mL tube after the tooth is removed, transfer this to the same 50mL conical tube, and mix gently. NOTE: We recommend using small, sterile forceps to remove each tooth from its 5mL tube by grasping the parafilm tail. It may also be possible to gently slide the tooth out of the 5mL tube by tilting the tube or using a sterile pipette tip. If this method is chosen, take care not to lose or contaminate any lysate that remains in the 5mL tube.
8. **Proceed with standard extraction cleanup:** Following a procedure based on Dabney et al. (2013) and Korlević et al. (2015), continue with the DNA purification.
    a. Transfer the binding buffer-lysate solution to the Roche High Pure Extender Assembly Tubes. Centrifuge the Roche High Pure Extender Assembly Tubes at 1,500 rpm for 4 minutes.
    b. Disassemble the High Pure Extender Assembly, seal and place the High Pure Spin Column in a 1.5 mL collection tube. NOTE: Be sure to label the spin column tubes.

c. Dry spin the spin column for 1 minute at 6000 rpm, to remove residual binding buffer.
d. Remove the spin column from the collection tube, discarding any flow through, and place the spin column in a fresh 1.5mL collection tube.
e. Add 650uL of PE wash buffer to the spin column, and centrifuge for 1 minute at 6,000 rpm. Discard the flow-through and repeat this washing step.
f. Place the spin column in a fresh collection tube and perform a dry spin at maximum speed for 1 minute to remove any residual wash buffer. Discard the collection tube and place the spin column in a fresh, labeled 1.5mL microcentrifuge tube.
g. Elute the DNA by adding 25uL of TET buffer to the silica membrane of the spin column. Incubate for 10 minutes at 37$^{o}$C. Centrifuge at maximum speed for 30 seconds. Repeat this step to yield a total of 50uL of DNA extract.

9. **Clean the tooth:** Remove the parafilm from the tooth, and rinse with abundant amounts of water or 95% ethanol in order to wash away any remaining extraction buffer from the tooth surface. Allow the tooth to air dry for a minimum of 24 hours. NOTE: Be extremely careful removing the Parafilm from teeth, as it is possible to break teeth if too much force is used. This is especially important for poorly preserved, low-density samples. It may help to use a sharp blade to cut the parafilm (Supplementary Figure 6C).
